# Molecular Detection of *H.pylori* Antibiotic-Resistant Genes and Bioinformatics Predictive Analysis

**DOI:** 10.1101/325654

**Authors:** Dan Wang, Qianqian Guo, Zhi Lv, Yuan Yuan, Yuehua Gong

## Abstract

To explore the mutation characteristics of *H.pylori* resistance-related genes to antibiotics of clarithromycin, levofloxacin and metronidazole. 23S rRNA, *gyrA, gyrB, rdxA* and *frxA* genes were amplified and sequenced, respectively. Their structural alteration after mutation was predicted using bioinformatics software. In the clarithromycin-resistant strains, the mutation rate in site A2143G was 74.2% (n=23). The mutations in sites C1883T, C2131T and T2179G might cause structural alteration. In the levofloxacin-resistant strains, the mutation rates in 87 (N to K/I) and 91 (D to N/Y/G) of *gyrA* were 28.6% (n=16) and 12.5% (n =7), respectively. Meanwhile, one of the mutation strains in site 91 was accompanied by D99N variation. Additionally, a D143E mutation was found in one drug-resistant strain. Some changes of tertiary structure occurred after these mutations. The mutation types of RdxA protein consisted of protein truncation caused by premature stop codons (n=26, 33.3%), frameshift mutations (n=8, 10.3%), FMN-binding sites (n=16, 20.5%) and the others (n=11, 14.1%). Predictive analysis showed that mutations in the first three groups and the A118S of the last group could lead to structural alteration. Our study suggested the clarithromycin-resistant sites of *H.pylori* were mainly located in A2143G of 23S rRNA. C1883T, C2131T and T2179G might also be related to resistance. Levofloxacin resistance was mainly based on the amino acid changes in 87 and 91 sites of *gyrA.* The new sites D99N and D143E might also be associated with resistance. Metronidazole resistance was related to RdxA protein truncation, frameshift, and FMN binding. The new site A118S might also be linked to drug resistance.

---

It has been well known that eradicating *H.pylori* can prevent the progression of diseases and reduce the risk of gastric cancer (1). Currently, the antibiotics used for *H.pylori* eradication are mainly composed of proton pump inhibitors and gastric mucosal protectant in combination with one or two kinds of antibiotics, including clarithromycin, metronidazole, levofloxacin, and amoxicillin, etc. With the extensive implementation of eradication therapy, the resistance of *H.pylori* to antibiotics has been increasing year by year (2).

At present, bacterial factors are regarded as the main cause for antibiotic resistance. Some changes of *H. pylori* genes could result in the loss of antibiotics targets and thus induce bacterial resistance. For example, it has been suggested that the A2143G, A2142G and A2142C mutations in 23S rRNA V region are related to clarithromycin resistance (3, 4). However, the mutation sites of drug resistance-related genes vary from place to place. For instance, *Kim* et al reported the resistant sites of *H.pylori* to clarithromycin were located at A2143G and T2182C in South Korea (5), while A2143G and A2146G existed in Nepal (6). A research in Brazil showed the resistance mutation was mainly located in A2147G (7). Other new mutation sites including T2115G, G2141A, T2190C, C2195T, A2223G, C2694A (8), C2147G, G1939A and T1942C (9) have also been mentioned in other areas. Moreover, the quinolone resistance determining region (QRDR) affected by *gyrA* and *gyrB* mutations of *H.pylori* has been shown to be associated with levofloxacin resistance. The main mutations occurred in amino acid sites of 91, 88 and 87 in *gyrA.* Among them, mutations in 87 and 91 were the most frequent, with the forms of N87 to I, K and D91 to Y, N, G (10, 11). With respect to metronidazole resistance, it was mainly related to the inactivation of oxidoreductase encoded by *rdxA.* The *rdxA* gene had diverse variation and no consistent site has been found yet. For *frxA,* some researches demonstrated it could increase the resistant level of metronidazole in the presence of *rdxA* mutations (12). However, it was also suggested that the frame shifting mutations of *frxA* existed both in susceptible and resistant strains, without association with metronidazole resistance (13). The above-mentioned reports indicate the relationship between mutation and drug resistance remains controversial.

Zhuanghe country, in Liaoning province of North China, is a high-risk region of gastric cancer. Previously, we detected the resistance phenotype of *H.pylori* strains there to clarithromycin, levofloxacin and metronidazole by E-test. In the present study, the *H.pylori* resistance-related genes were amplified and sequenced. Furthermore, their structural alteration was predicted using bioinformatics software. Our study aims to investigate the mutation characteristics of *H.pylori* resistance-related genes in the high-risk area of gastric cancer and provide theoretical basis for the exploration in molecular mechanisms of antibiotics resistance.

## RESULTS

### Molecular detection and bioinformatics predictive analysis of *H.pylori* resistance genes to clarithromycin

Considering 26695 strain as the reference, the 23S rRNA gene sequence was compared between 9 clarithromycin-sensitive and 31 clarithromycin-resistant strains. A2143G mutation occurred in 74.2% (n=23) resistant strains. In addition, some sporadic changes were also found, including C1883T, G1949A, C2131T, C1883T, G1949A, C1953T, T2179G, A2210C and G2211T (Figure 1). For the 2182 site previously reported to be related to clarithromycin resistance (14), T to C (n=28, 90.3%) or A (n=2, 6.5%) mutation existed in resistant strains. All the 9 (100%) sensitive strains had T2182C mutation.

**Figure 1.**
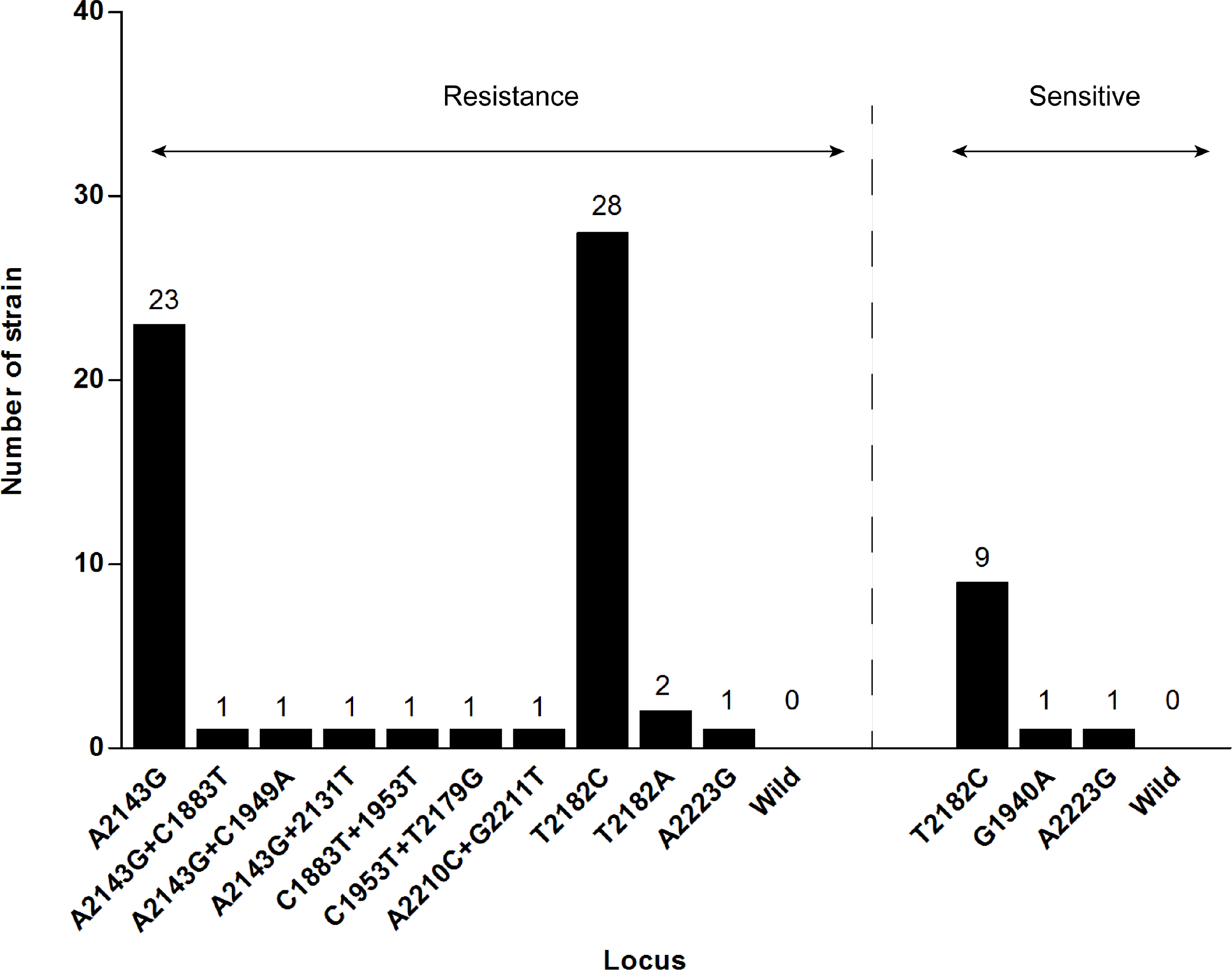
The distribution of mutatnt sites in the 23S rRNA V region of 9 clarithromycin sensitive and 31 resistant strains of *H.pylori.*

Next, Mfold software was used to predict the secondary structure of 23S rRNA. The base C of pre-mutation RNA was inside the loop, while the loop became narrowed and the adjacent helical region got longer after C1883T mutation (Figure 2). The C2131T mutation made the loop smaller, the helical region longer and the base T become part of it (Figure 3). After T2179G mutation, the hydrogen bonds disappeared. No structural alteration was found in the mutation sites of A2143G, T2182C/A, G1949A, G1953T, A2210C and G2211T.

**Figure 2.**
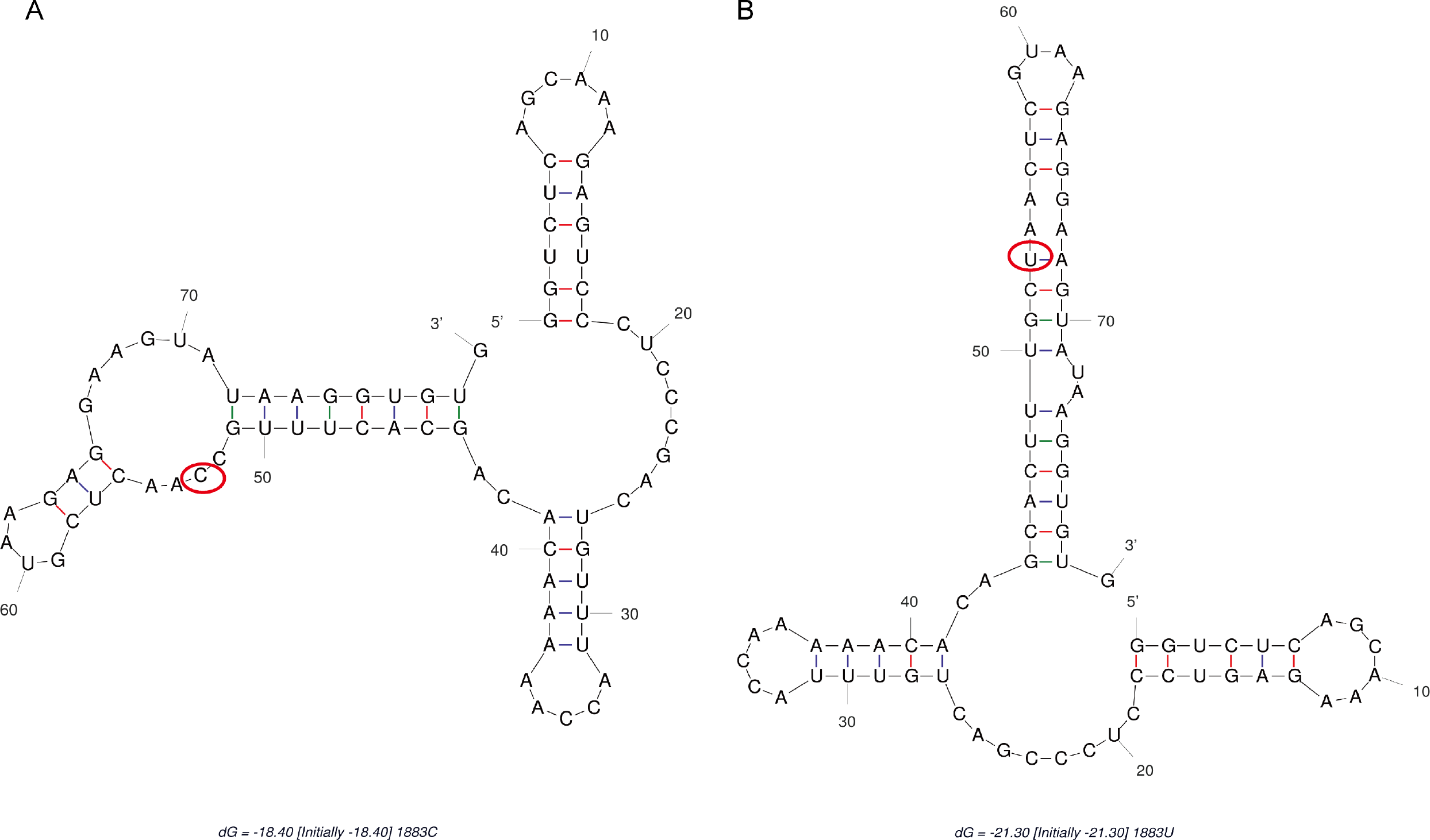
The prediction of RNA secondary structure before and after mutation of C1883T (U) in 23S rRNA V region by Mfold software. A. The base C of pre-mutation RNA secondary structure was inside the loop, B. after the C1883T mutation, the loop became narrowed and the adjacent helical region got longer.

**Figure 3.**
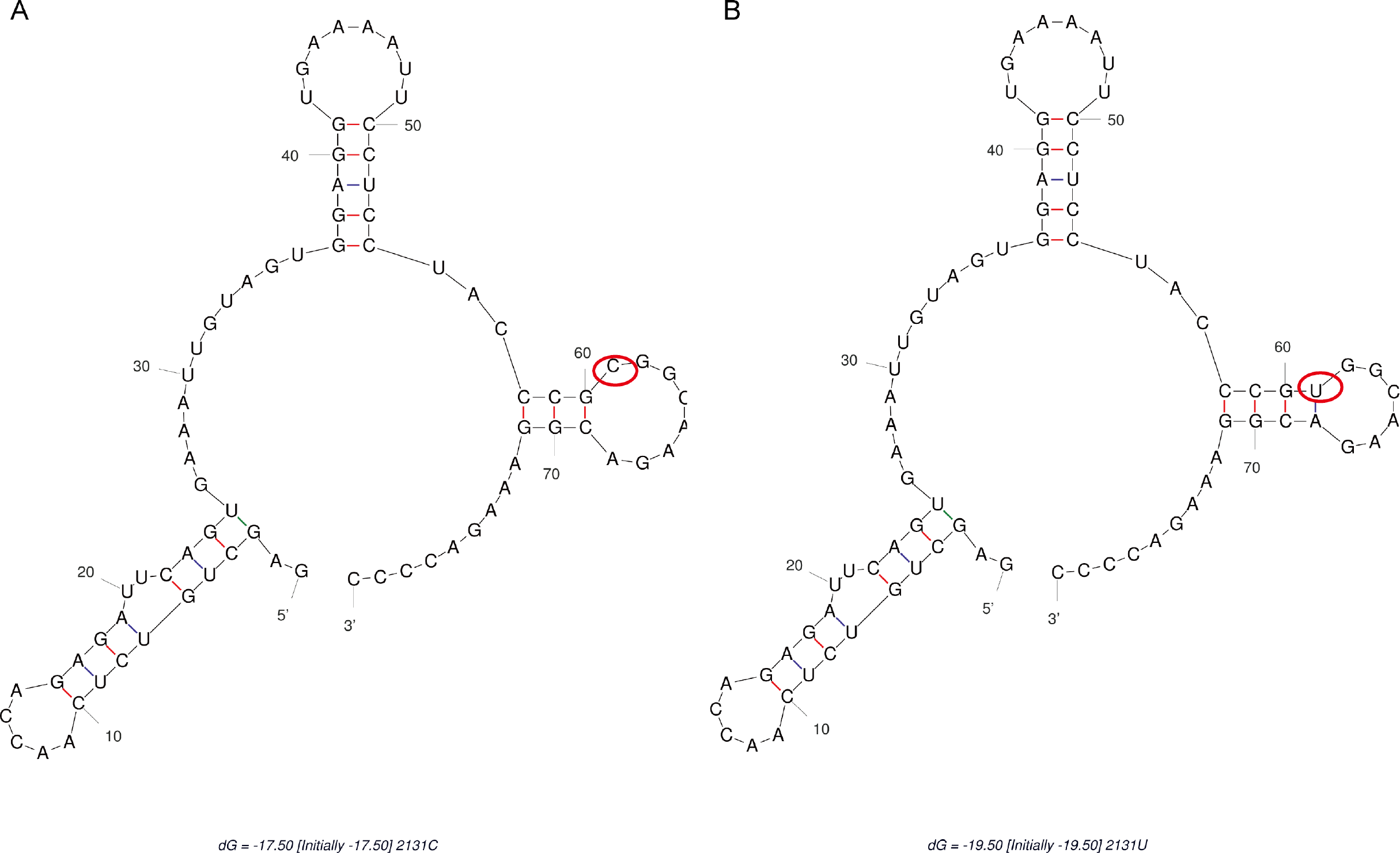
The prediction of RNA secondary structure before and after mutation of C2131T (U) site in 23S rRNA V region by Mfold software. A. The base C of pre-mutation RNA secondary structure was in the interior of the loop, B. after the C2131T mutation, the loop becomes smaller, the helical region longer and the base T become part of it

### Molecular detection and bioinformatics predictive analysis of *H.pylori* resistance genes to levofloxacin

The differences between 10 levofloxacin-sensitive and 56 levofloxacin-resistant strains were further compared when 26695 strain was taken as the reference. The mutation rates of 87 (N to K/I) and 91 (D to N/Y/G) in resistant strains were 28.6% (n =16) and 12.5% (n=7), respectively. Meanwhile, one of the mutation strains in site 91 was accompanied by D99N variation. Additionally, a D143E mutation was found in one resistant strain. No change of GyrA protein was observed in another 29 resistant strains (51.8%). In all the 56 resistant strains, the MIC values of mutant strains were significantly higher than those of non-mutant strains (30.73 ± 6.23 ug/L *vs* 7.70±11.94 ug/L, *P*< 0.001). We also detected the amino acid mutations of D481E and R484K for GyrB protein. However, the two mutations were discovered both in resistant (n=8, 14.3%) and sensitive strains (n=2, 20%). 48 resistant strains (85.7%) had no alteration in GyrB protein (Figure 4).

**Figure 4.**
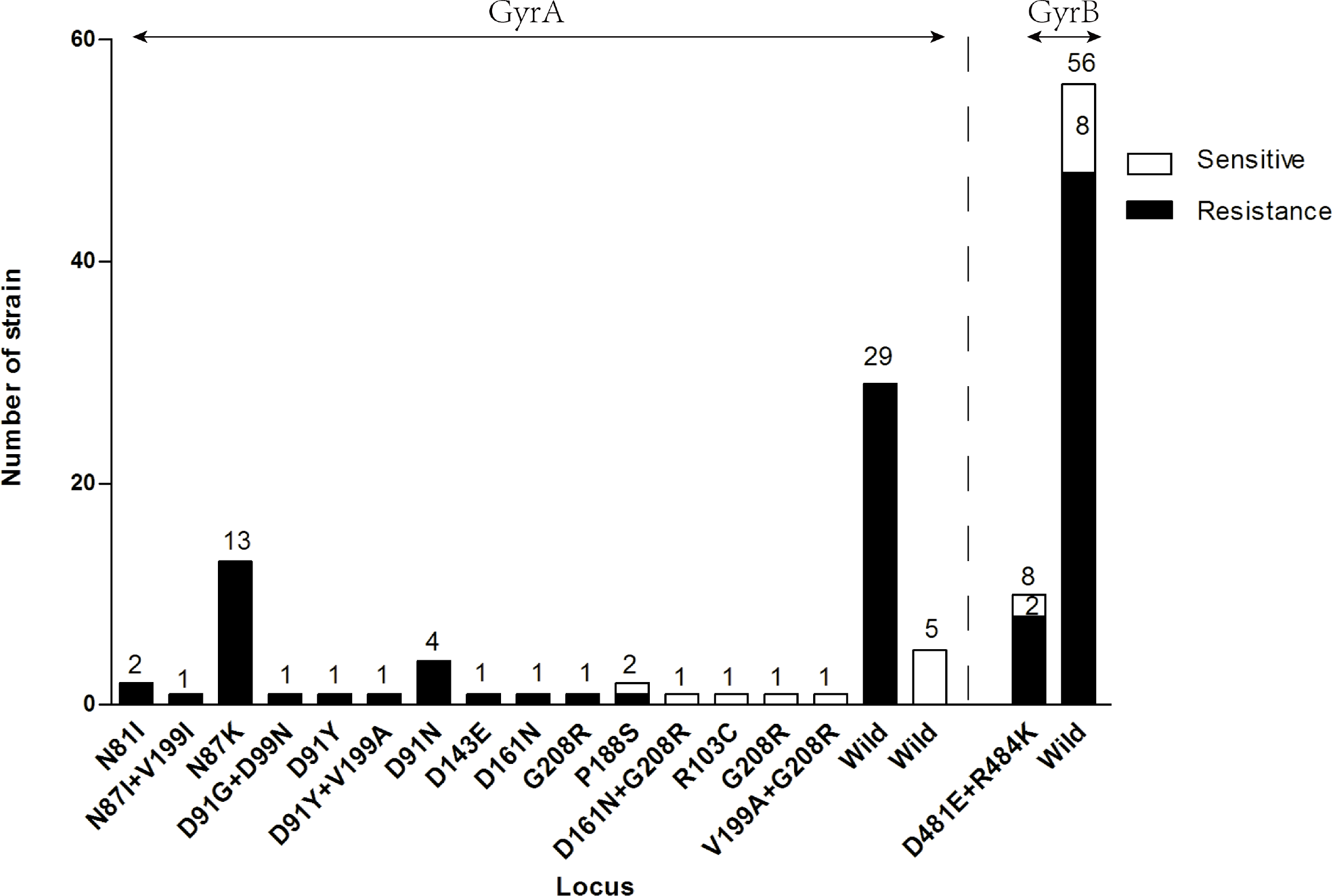
The distribution of GyrA and GyrB protein mutant sites in 10 levofloxacin-sensitive strains and 56 resistant strains of *H.pylori.*

Then the SWISS-MODEL software was used to model protein tertiary structure before and after mutation. The overlap matching of tertiary structure before and after N87I, D91Y/G/N and D143E mutations in GyrA protein was shown in Figure 5 A. The protein structure changed from partial fold to random curl after mutation. Moreover, the overlap matching of tertiary structure before and after N87K and D99N mutations was presented in Figure 5 B and C. The protein structure changed in both folded and irregularly curled region after mutation. However, the D481E and R484K mutations did not cause significant alteration in GyrB protein (Figure 5 D & E).

**Figure 5.**
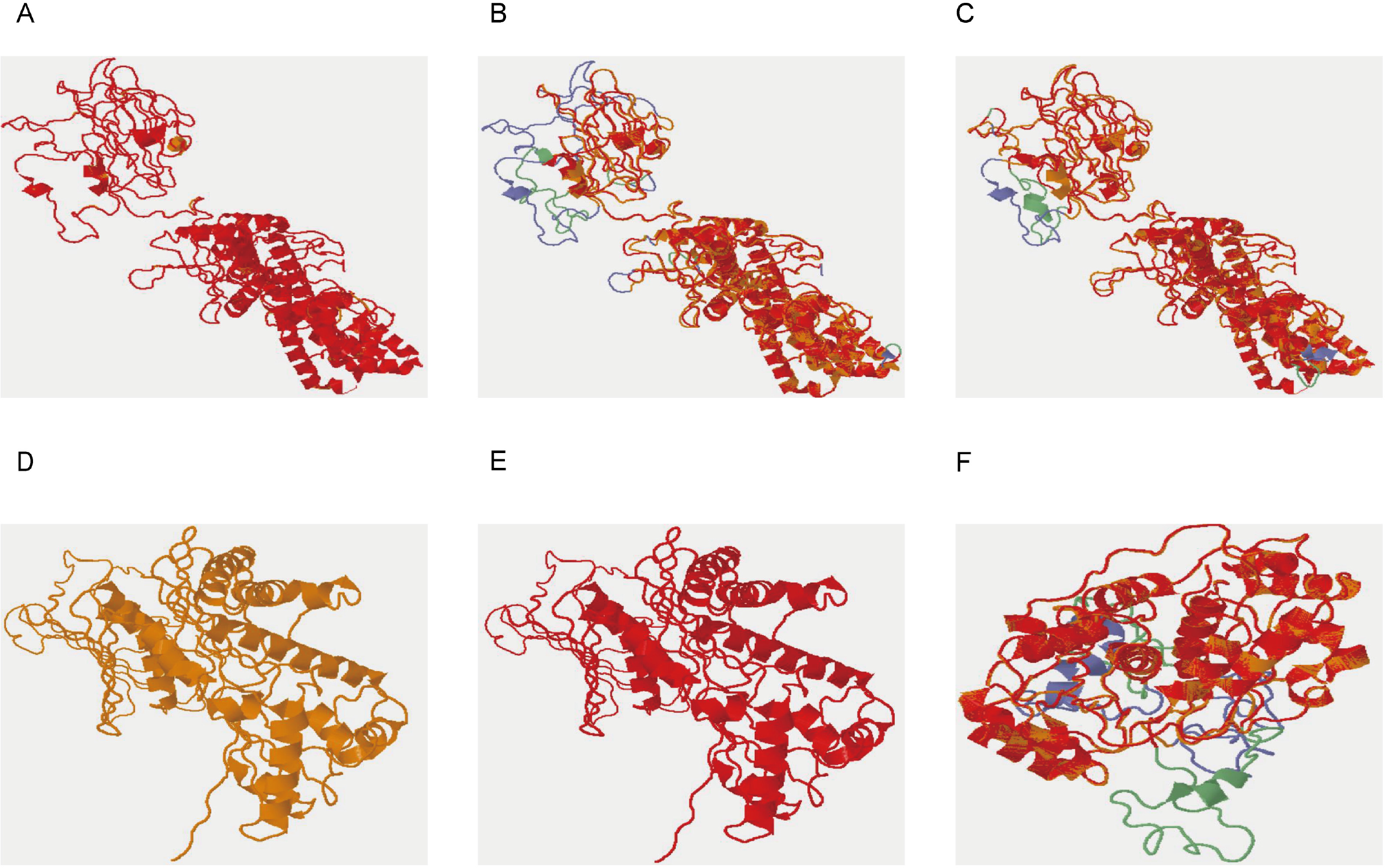
Prediction of tertiary structure before and after mutation of GyrA, GyrB and RdxA protein site by SWISS-MODEL software. The yellow indicates the tertiary structure of the protein constructed using the pre-mutated sequence, the red indicates the tertiary structure of the corresponding protein constructed by using the mutated sequence, and the purple and green regions represent the differences between the two protein structures before and after the mutation. The purple is an un-mutated structure, green is the mutated structure. A: The overlap matching of tertiary structure before and after N87I, D91Y / G / N and D143E mutations in GyrA protein, the protein structure changed from partial fold to random curl after mutation. B, C: The overlap matching of tertiary structure before and after N87K and D99N mutation of GyrA protein, the protein structure changed in both folded and irregularly curled regions after mutation. D, E: D481E and R484K of GyrB protein before and after mutation were completely consistent before and after the mutation, and no obvious change was found. F: RdxA protein A118S mutation before and after the tertiary structure of the overlapping match, the structure in 100-127 region of A chain changed from regional fold into random curl, and partial fold appeared in 91-130 region of B chain with random curl after A118S mutation.

### Molecular detection and bioinformatics predictive analysis of *H.pylori* resistance genes to metronidazole

With the reference of 26695 strain, the differences between 9 metronidazole-sensitive strains and 78 metronidazole-resistant strains were also investigated. RdxA protein mutations were suggested to be random, wide and scattered, without obvious regular pattern. Compared with the susceptible strains, RdxA protein mutations were classified into four types. First, 26 (33.3%) strains had protein truncation due to premature stop codons, mainly including amino acid sites of 33, 35, 50, 65, 73, 76 and 113, etc. Second, 8 (10.3%) strains had frameshift mutations caused by base insertion or deletion, mainly including 30, 39, 58, 188, 189 and 192 sites. Third, 16 (20.5%) strains had mutations in flavin-mononucleotide (FMN) binding site, mainly including R16H/C, S45N, S46F, Y47C, N48Y, I160V, G162R, and G163D. Fourth, 11 (14.1%) strains had other highly frequent mutations (cases>3), mainly including M21V/I/A, T58A, K64N, A68T/V, A118S and G150S (Table 1). Besides, no genetic variation was found in 17 (21.79%) resistant strains. Among the 78 drug-resistant strains, the difference of MIC values with or without mutation did not reach statistical significance (39.79ug/L *vs* 38.47ug/L, *P*=0.744).

**Table 1.**
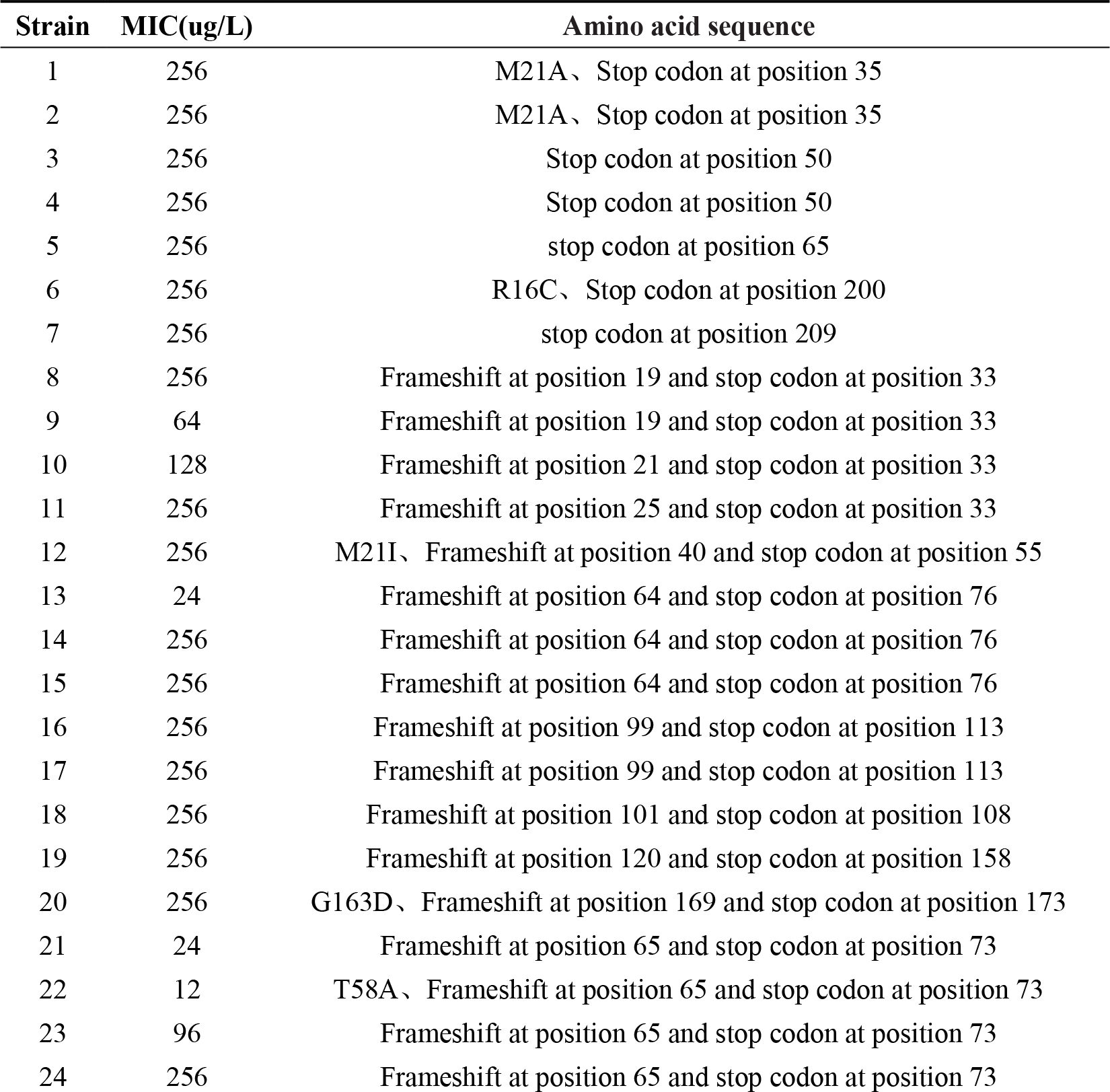
Mutational events in metronidazole resistance strains.

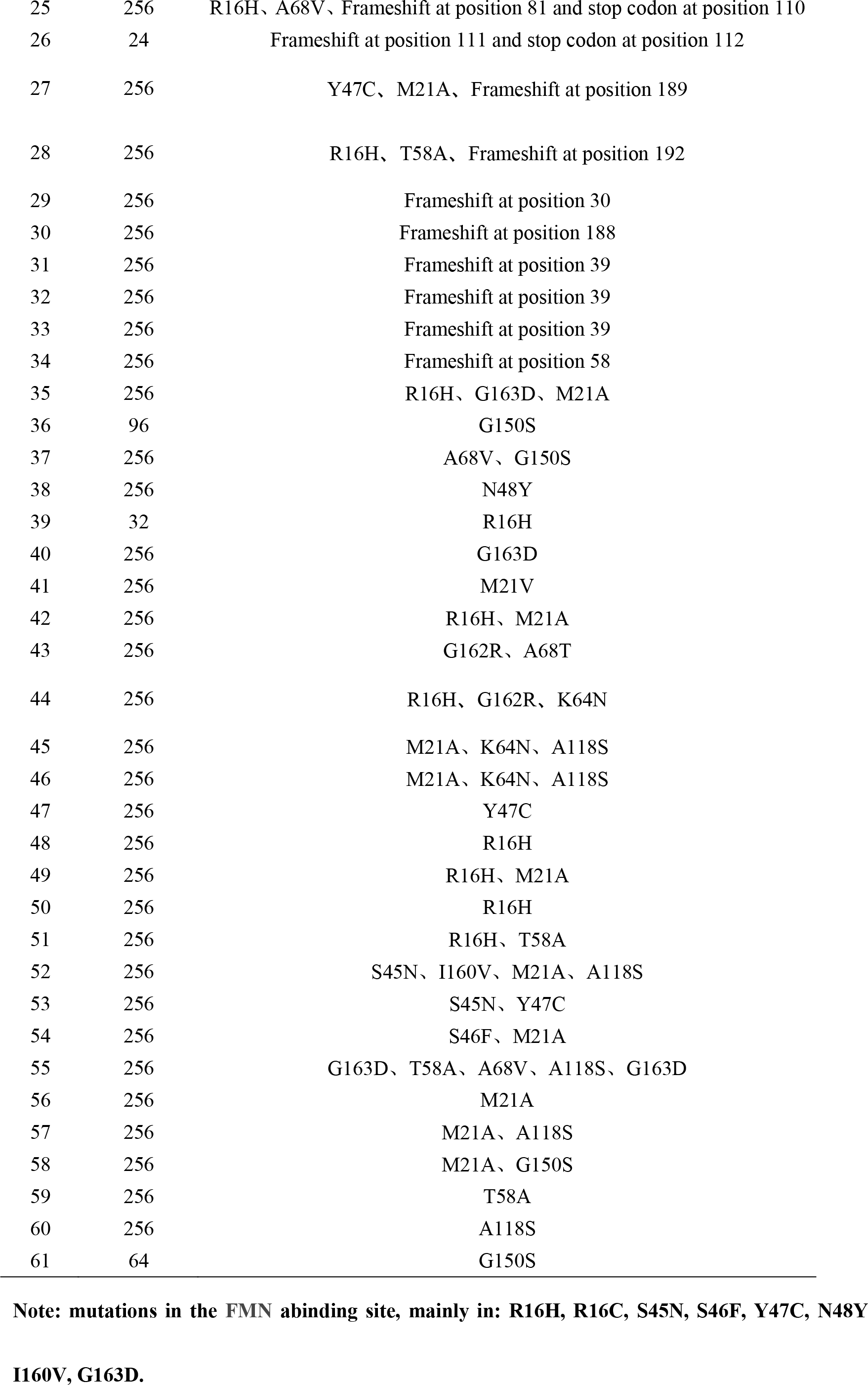

Similarly, we compared the amino acid sequence of FrxA protein between 9 sensitive and 78 resistant strains with the reference of 26695 strain. The distribution of mutation sites was also shown to be random, wide and scattered, without regular pattern. Compared with the sensitive strains, FrxA protein mutations could also be categorized into four types (Supplemental Table 2). In the resistant strains with RdxA mutation, whether FrxA protein mutation was present or not made no effect on *H.pylori* resistance to metronidazole (211.00 ± 89.76 ug/L *vs* 233.70 ± 61.70 ug/L, *P*>0.05).

Furthermore, the tertiary structure of RdxA protein before and after mutation was predicted using the SWISS-MODEL software. The first three types of protein mutation mentioned above could cause structural alteration (data not shown). As for the fourth one, we found the structure in 100-127 region of A chain changed from regional fold into random curl, and partial fold appeared in 91-130 region of B chain with random curl after A118S mutation (Figure 5 F).

## DISCUSSIONS

Antibiotic resistance is the leading cause for the failure of *H.pylori* eradication. Previously, high resistance rates of *H.pylori* to clarithromycin, levofloxacin, and metronidazole were found in the high-risk region of gastric cancer by our research group. In the present study, mutations of *H.pylori* resistance-related genes to the three antibiotics were identified, and structural alteration was further predicted using bioinformatics software, aiming to explore the molecular mechanisms associated with *H. pylori* resistance.

Clarithromycin is a first-line drug for *H.pylori* eradication and drug resistance is the main cause for poor curative effect. Lots of researches have revealed the most common mechanism of *H. pylori* resistance to clarithromycin, which is the mutation of A2143G, A2142G/C and A2144G located in 23S rRNA V region (8, 9, 15, 16). In our study, we found the main resistant mutation of *H.pylori* to clarithromycin was A2143G. Moreover, the C1883T, G1949A, G1953T, C2131T, T2179G, A2210C and G2211T sites were newly discovered in resistant strains, which had not been reported yet. Structure prediction showed the C1883T, C2131T, and T2179G mutations could alter the secondary structure of 23S rRNA. Therefore, it can be inferred that these 4 sites may be related to clarithromycin resistance. In addition, A2143G was well accepted to be associated with the resistance. However, we found the A2143G mutation could not lead to structural alteration, suggesting that it might not affect RNA conformation but clarithromycin binding. Although T2182C/A was indicated to be associated with resistance by previous study (17), it was observed both in resistant and susceptible strains without structural alteration. *Burucoa* et al performed natural transformation for T2182C in *H. pylori* and found genetic variation did not affect clarithromycin resistance (18). In conclusion, the mutation in site 2182 may be irrelevant to clarithromycin resistance.

It is generally believed that mutations in the amino acid 91, 87, and 88 sites of *H. pylori gyrA* are involved in levofloxacin resistance (11, 19). Our study suggested the main resistance-related sites are 87 and 91, and the new sites 99 and 143 might also be linked to drug resistance. Prediction analysis showed the structure in 87 and 91 significantly changed after mutation. *Matsuzaki* et al found amino acid 87 and 91 sites were located in the N-terminal region of GyrA protein combined with DNA (20). Therefore, the 87 and 91 mutations might not only affect protein structure but also directly impair the binding between protein and DNA. Regarding 99 and 143 sites, bioinformatics prediction demonstrated the protein structure changed from fold to random curl after mutation. The two sites may be related to levofloxacin resistance, which needs verification by further investigations. Additionally, no *gyrA* change was shown in 29 (51.8%) resistant strains. *Seck* et al also found no *gyrA* mutation in some levofloxacin-resistant strains (21), suggesting other resistance-related mechanisms such as efflux pump, membrane permeability, autophagy, globular transformation and even new resistant genes, etc (22). The MIC values of drug-resistant strains with *gyrA* mutation were significantly higher than those of non-mutant strains, indicating that *gyrA* mutation was associated with high levofloxacin resistant level. With respect to *gyrB* gene, previous research revealed the variation in amino acid D481E and R484K sites were related to levofloxacin resistance (23), and could further promote levofloxacin resistance under *gyrA* mutation (24). However, we found D481E and R484K were present both in drug-resistant and susceptible strains without increasing MIC level of *H.pylori* resistance to levofloxacin. And bioinformatics predictive analysis also showed no effect of mutations in GyrB 481 and 484 on the tertiary structure. As a result, they might have no association with drug resistance.

It has been well acknowledged that oxidoreductase inactivation is the main cause for *H.pylori* resistance to metronidazole, which is induced by frame mutation, insertion or deletion mutations in *rdxA* and *frxA* genes. Our study demonstrated the distribution of RdxA and FrxA mutation sites was random, scattered, and irregular. They were classified into four types, including protein truncation due to premature stop codons, frameshift mutations, FMN-binding sites, and other highly frequent mutation sites. These variations have also been reported in other region. For example, *Kwon* et al found frameshift mutation in metronidazole-resistant strains leading to early termination of protein translation (25). A mutation in site 16 binding to FMN was discovered in resistant strains by *Secka* et al (26). Bioinformatics predictive analysis showed the protein structure changed significantly after the first two categories of mutation. In the third group, the structural alteration of protein occurred in R16H/C, S45N, S46F, Y47C, N48Y, I160V, G162R, and G163D. Most scholars believed that transformation in the sites binding to FMN could impair the non-covalent combination with RdxA protein and thus result in obstruction of electron transport (26). Therefore, they are likely to be associated with metronidazole resistance. Among the fourth type of highly frequent mutation sites including M21V/I/A, T58A, K64N, A68T/V, A118S and G150S, only the A118S site was shown to induce structural alteration. And A118S/T has been reported to be closely related to metronidazole resistance (27, 28). Additionally, some drug-resistant strains have no *rdxA* mutation, suggesting other resistant mechanisms such as *rpsU* mutation other than *rdxA* (12), which needs further investigation.

It remains controversial about the relationship between *frxA* and metronidazole resistance. Some scholars hold that *frxA* variation might be involved in high MIC level (29), while others believed it had nothing to do with metronidazole resistance (13). Our study showed the protein translation truncation of FrxA due to premature stop codons occurred both in resistant and susceptible strains. However, no change of RdxA protein was found in them. Besides, *frxA* could not enhance metronidazole resistance, as a result of which, *frxA* might not affect the sensitivity of *H.pylori* to metronidazole.

In summary, we detected the mutation sites of *H. pylori* to three antibiotics and further performed prediction analysis using bioinformatics software. The results showed the clarithromycin-resistant sites of *H.pylori* were mainly located in A2143G of 23S rRNA. C1883T, C2131T and T2179G might also be related to drug resistance, while T2182C showed no association. Levofloxacin resistance was mainly based on the amino acid changes in 87 and 91 sites of *gyrA.* The new sites D99N and D143E might also be associated with it, while the *gyrB* gene had no relationship with bacterial drug resistance. Metronidazole resistance was related to RdxA protein truncation, frameshift and FMN binding. The new site A118S might be linked to drug resistance, while *frxA* did not contribute to it. The study would provide novel ideas and methods for the identification of *H.pylori* resistance-related mutations, and also clues and basis for further investigation on specific molecular mechanism. In addition, many resistant strains were found to have no variation in the above-mentioned genes. That indicates the complex mechanism of *H.pylori* resistance, and other possibilities should also be taken into account such as membrane permeability, efflux pump or new genes related to drug resistance, which needs further in-depth exploration.

## MATERIALS AND METHODS

### H.pylori strains

A total of 9 clarithromycin-sensitive and 31 clarithromycin-resistant strains, 10 levofloxacin-sensitive and 56 levofloxacin-resistant strains, 9 metronidazole-sensitive and 78 metronidazole-resistant strains were involved in our study. The study was approved by the Ethics Committee of China Medical University and informed consents were obtained from all the participants.

### DNA extraction

The whole genomic DNA of *H. pylori* was extracted using phenol-chloroform method and the details were described in a previous study by *Gong* et al (30). The DNA was dissolved in 50-100 μl TE buffer. Ultraviolet spectrophotometer (Thermo, Germany) was used to measure the concentration and purity of DNA. Then the specimen was kept in −20°C to preserve.

### Polymerase chain reaction and DNA sequencing

Polymerase chain reaction was used to amplify the domain V of 23S rRNA, *gyrA* (hp0701), *gyrB* (hp0501), *rdxA* (hp0954) and *frxA* (hp0642). The amplification system was comprised of 2 mmol/L dNTP, 2.5 U Taq or LA Taq DNA polymerase, 10 × buffer or LA Buffer II (Takara, China), 10 pmol primer, and 2μl of DNA template. 2x Taq PCR StarMix Kit (GeneStar, China) was used for the samples failing to be amplified without changing the amounts of primer and template. The reaction conditions were set as the followings: denaturation at 94 °C for 1 min, annealing shown in Supplemental Table 1 and extension at 72 °C for 1 min. Primer synthesis and sequencing of amplified products were completed by The Beijing Genomics Institute (BGI).

### DNA Sequence Analysis

The NCBI website (http://blast.ncbi.nlm.nih.gov/Blast.cgi) was used to determine sequencing accuracy by referring to *H.pylori* 26695 (GenBank accession number AE000511.1 GI: 6626253). The DNAStar (version 7; Lasergene, USA) and DANMAN (version 6; DNAMAN, USA) softwares were used to edit, align, translate and analyze the gene or amino acid sequences. Secondary structure prediction of 23S rRNA before and after mutation was performed using Mfold (http://www.bioinfo.rpi.edu/applications/mfold). *E.coli* (PBD ID: 1ZVU and 4JUO) was chosen as a reference for GyrA and GyrB protein, and *H.pylori* (PDB ID:3QDL) was the reference for RdxA protein. SWISS-MODEL (https://swissmodel.expasy.org/) was adopted to simulate the tertiary structure of GyrA, GyrB and RdxA protein before and after mutation. TopMatch (https://topmatch.services.came.sbg.ac.at/) was used for online comparison.

### Statistical analysis

All statistical analyses were conducted by the SPSS statistical software package version 18.0 (Chicago, IL, USA). Mann-Whitney *U* or t-tests were used to test the differences in Minimum inhibitory concentration (MIC) value between the mutants and non-mutants in the resistant strains. *P* value less than 0.05.was accepted to be statistically significant.

